# Stochastic Spatiotemporal Simulation of a General Reaction System

**DOI:** 10.1101/2022.10.26.512711

**Authors:** Andrew J. Loza, Marc S. Sherman

**Author notes:** Corresponding author *Email address:* (Andrew J. Loza).

## Abstract

Biological systems frequently contain biochemical species present as small numbers of slowly diffusing molecules, leading to fluctuations that invalidate deterministic analyses of system dynamics. The development of mathematical tools that account for the spatial distribution and discrete number of reacting molecules is vital for understanding cellular behavior and engineering biological circuits. Here we present an algorithm for an event-driven stochastic spatiotemporal simulation of a general reaction process that bridges well-mixed and unmixed systems. The algorithm is based on time-varying particle probability density functions whose overlap in time and space is proportional to reactive propensity. We show this to be mathematically equivalent to the Gillespie algorithm in the specific case of fast diffusion. We develop a computational implementation of this algorithm and provide a Fourier transformation-based approach which allows for near constant computational complexity with respect to the number of individual particles of a given species. To test this simulation method, we examine reaction and diffusion limited regimes of a bimolecular association-dissociation reaction. In the reaction limited regime where mixing occurs between individual reactions, equilibrium numbers of components match the expected values from mean field methods. In the diffusion limited regime, however, spatial correlations between newly dissociated species persist, leading to rebinding events and a shift the in the observed molecular counts. In the final part of this work, we examine how changes in enzyme efficiency can emerge from changes in diffusive mobility alone, as may result from protein complex formation.

## 1. Introduction

Biological processes are often characterized by slowly diffusing components present in small numbers, as well as reactions that depend nonlinearly on substrate concentrations and may not be at equilibrium [1, 2, 3]. In this realm, fluctuations in the number and spatial distribution of the participating species become non-trivial and can lead to functional consequences [4]. This variation has been demonstrated to play a role in a wide variety of cellular processes, including fate in bacterial populations following viral infection or nutrient restriction [5, 6], phenotypic variation in genetically identical populations [7, 8, 9], and cell differentiation in organism development [10].

Mathematical models are critical for understanding and predicting the behavior of biological systems, but deterministic models often break down in describing biological dynamics [11]. Molecular concentration becomes discontinuous in the spatial domain, due to slow diffusion, and in magnitude, due to countable fluctuations in the number of molecules. To understand the details of when and how fluctuations influence the behavior of biological systems, a variety of mathematical models have been developed to capture the discrete and stochastic nature of the systems involved [12, 13, 14, 15, 16, 17]. A computational approach was provided by Gillespie to simulate the chemical master equation [12]. In the original work, spatial distribution of molecules was assumed to be uniform, but an extension of the model using subvolumes allowed for simulation of spatial heterogeneity provided each subvolume fulfilled the well-mixed requirement [18, 14, 13]. Related work divided space into metastable compartments, simulating reactions within compartments and transitions between them [19]. A second class of simulation techniques relies on a molecular dynamics type approach, simulating the diffusive trajectories of species involved [15]. For simulations when the diffusive steps are small compared to inter-particle distance, this approach can become inefficient. To remedy this, methods referred to as Green’s Function Reaction Dynamics (GFRD) and First-Passage Kinetic Monte Carlo (FPKMC) provided a strategy to jump forward in time to when interactions occur [16, 20, 17].

While these stochastic and discrete models extend deterministic and continuous approaches, they retain a few disadvantages. Subvolume methods may be influenced by the selection of subvolume size, with potential inaccurate results [21], and GFRD and FPKMC require careful treatment of certain spatial arrangements of molecules that, if not explicitly accounted for, can limit computational efficiency [17, 22, 23, 24]. Furthermore, processes that span timescales and length-scales may be coupled in biological systems, complicating the choices required to efficiently and accurately implement these algorithms. Therefore, further development of methods that address these shortcomings is warranted to advance the field.

Here we present the framework and implementation of an event driven stochastic spatiotemporal simulation of a generalized reaction process which bridges diffusion-limited stochastic kinetics with well-mixed reaction kinetics. The analytic framework of this algorithm is unique in its ability to span well-mixed and non-mixed systems, providing a seamless way to simulate complex biological processes. In the first part of this work, we develop the underlying reaction probability density function and the sampling procedure to determine reaction type, location, and participating molecules.

We additionally demonstrate that this method is mathematically equivalent to the Gillespie algorithm in the specific limit of rapid diffusion. In the second part of this work, we provide an implementation for a bimolecular association-dissociation reaction and demonstrate the necessity for reaction-diffusion modeling in predicting expected values when re-binding probabilities are high. We then provide strategies for increasing computational efficiency for systems with large numbers of particles. Lastly, we examine the influence of protein complex formation on enzyme efficiency solely through changes in diffusive mobility. This framework relies on general particle probability density functions without making assumptions about their forms. Here we focus on diffusive motion, however this framework could be used to simulate systems with more complex processes driving particle mobility.

## 2. Simulation Algorithm

### 2.1. System Definitions

Let the general system be defined as a set of *M* mobile species *S*_*i*_ that may interconvert through *N* reactions *R*_*µ*_ within a volume *V*. We are interested in systems with a discrete number of members of each species and therefore define *X*_*i*_ as the current number of molecules of *S*_*i*_. Individual particle identity is accounted for through a second subscript *j* such that *S*_*ij*_ refers to the *j*^*th*^ molecule of the *i*^*th*^ species, where 1 ≤ *j* ≤ *X*_*i*_.

### 2.2. Derivation of the Reaction Probability Density Function

The Gillespie algorithm requires the computation of the reaction probability density function, *P*(*τ, µ*), defined as the probability at time *τ* that the next reaction in *V* will occur in the differential time interval *δτ* and will be of type *µ* [12]. The goal of the present work is to extend this joint probability density function to include location and molecular identity by deriving 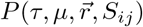: The probability at time *τ* that the next reaction in V will occur in the differential interval *δτ*; *and* will be of type *µ*; *and* will occur at position 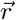; *and* will involve the specific molecules *S*_*ij*_.

We first derive the univariate distribution *P*(*τ*) and then decompose it into the multivariate distribution 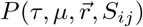, which can be sampled using Monte Carlo methods. Here we focus on a bimolecular reaction between two distinct species, but the derivation can be generalized for multiple reaction types. The reaction is:

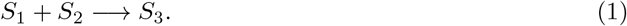

Two molecules have the potential to react if they collide with proper orientation and energy. A collision will take place whenever the centers of two spherical molecules come closer than:

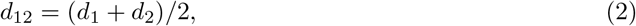

Where *d*_12_ is the collision distance threshold, *d*_1_ is the diameter of an *S*_1_ molecule, and *d*_2_ is the diameter of an *S*_2_ molecule. If the average speed of an *S*_1_ molecule relative to an *S*_2_ molecule is ⟨*v*_12_⟩, the *S*_1_ molecule produces a potential collision volume over time that can be defined as:

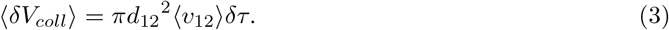

By determining the probability that this collision volume is occupied by an *S*_2_ molecule, the likelihood that the collision was successful, and the number of potential reactive collisions, Gillespie defines the total probability that a reaction *R*_*µ*_ will occur in *V* in the next time interval *δτ* as *h*_*µ*_*c*_*µ*_*δτ*, where *h*_*µ*_ is the number of distinct combinations of reactant molecules and *c*_*µ*_ is the probability that a particular combination of reactant molecules will indeed react [12]. The present goal is to determine the value of *h*_*µ*_*c*_*µ*_*δτ* without making the assumption of spatial homogeneity. To begin, we define *z*_*µ*_(*τ*)*δτ* as the probability that the next reaction *R*_*µ*_ will occur in *V* in the next time interval *δτ* beginning at time *τ*. Importantly, while *h*_*µ*_*c*_*µ*_ is constant with respect to time for any inter-reaction interval, *z*_*µ*_ may be a function of time.

A volume *V* can be broken into an infinite number of small sub-volumes in which the next reaction could occur. Because the probability of two reactions occurring in the time interval *δτ* vanishes as *δτ* goes to zero, the probabilities of this single next reaction being in each sub-volume are disjoint. This means, through the addition rule of disjoint events, that the sum of probabilities in all these sub-volumes is equal to *z*_*µ*_*δt*. In the limit of infinite sub-volumes, summation becomes integration, and we define a new variable, 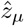:

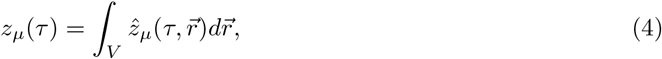

which is the probability density of the next reaction occurring at a specific position 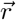 in the differential interval beginning at time *τ*.

In a general sense, 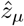 describes a conditional probability, namely:

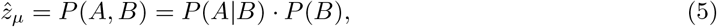

Where event *A* is an *R*_*µ*_ reaction and event *B* is the presence of an *S*_1_ molecule at a location. Computation of *P*(*B*) will be addressed first, followed by *P*(*A*|*B*).

The term *P*(*B*), or the probability that an *S*_1_ molecule is at a specific location, can be defined as follows. If the location of a molecule is known at a time *t* = 0, its location at *t* ≥ 0 is unknown precisely, but it is describable in terms of probability if the driving forces behind motion are known. Here we assume that thermal fluctuations drive molecular motion, yielding a Brownian diffusion process, however this method can be generalized to any source of motion that can be described in terms of a probability density function for each member of each species: 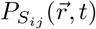. When multiple *S*_*i*_ molecules are present, a pseudo-probability density function describing probabilities of finding *S*_*i*_ at a point can be written:

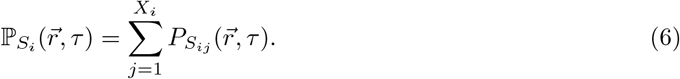

This is not a true probability density function because while the different 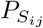 are independent, they are not disjoint. Therefore, the union of the individual 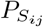 probability distributions is not simply the sum of probabilities of individual events but rather must be adjusted by the intersection of these probabilities. However, the cost of neglecting the intersection terms depends on the absolute probabilities of events, and as the absolute probabilities of the distinct events approach zero, the fractional error from neglecting their intersection becomes negligible (the probability of individual events is O(*δ*), the probability of intersection events is O(*δ*^2^) and higher). Therefore, if we consider a region where all the individual 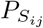 are high and query the probability that any *S*_*i*_ molecules are in that region, the integration of 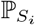 over the spatial limits of that region will be a poor approximation of the true probability (integrating 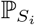 over *V* makes this readily apparent, as a value greater than 1 is seen). If, however, the region of interest is very small, such as *δV*_*coll*_, then the probability of individual molecules being within the region is small (computed by integrating 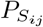 over the limits of the region *δV*_*coll*_), and as *δτ* drops to zero, the probability of two particles being within the collision volume is negligible. Thus, *P*(*B*) in (5), or the probability that an *S*_1_ molecule is within an infinitesimal volume in space, can be well approximated by 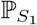 :

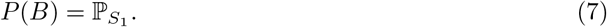

The first term on the right hand side of (5), *P*(*A*|*B*), is the probability that a reaction will occur given there is an *S*_1_ molecule at a specific location. Following the logic used by Gillespie, we break this into the probability that an *S*_2_ molecule is within the collision volume of a given *S*_1_ molecule and the probability that such a collision is productive. Using *θ*_*R*_ as the probability a collision is productive, *P*(*A*|*B*) becomes:

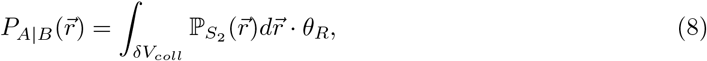

Where 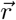 is the location of a given *S*_1_ molecule and *δV*_*coll*_ is the collision volume as defined in (3). The sum rule approximation employed here by the use of 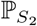 is valid because the collision volume is vanishingly small. The probability of any single molecule being within it is small, so any higher order terms that result from the product of individual probabilities will be negligible. To simplify this expression, we recognize that *δV*_*coll*_ is small such that 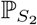 is essentially constant throughout this domain. Thus (8) becomes:

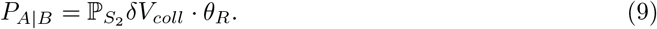

We now can re-write (5) substituting in (3), (7), (9):

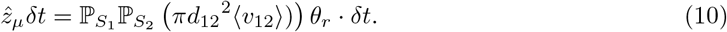

Integrating 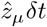 with respect to its spatial domain over a volume *V* gives the probability that a reaction occurs in the volume, *z*_*µ*_*δt*, as defined in (4). The only terms that depend on space are the pseudo-probability distributions, 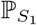 and 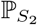, allowing for simplification:

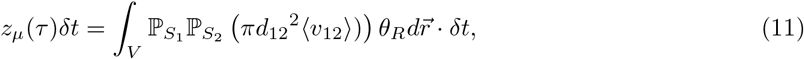

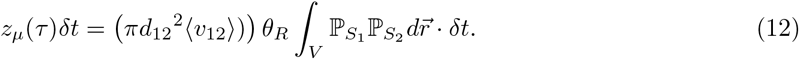

Defining the quantity of *z*_*µ*_*δt* allows us to determine the reaction probability distribution function *P*(*τ, µ*), the probability that the next reaction occurs at time *τ* and is a *µ* reaction. This probability can be decomposed into two components. The first component is *P*_0_(*τ*), the probability that no reaction occurs over time *τ*. The second component is *z*_*µ*_(*τ*)*dτ*, the probability that a reaction occurs in an infinitesimal window right after *τ* and has already been defined in (12). Because reactions are independent, random events that occur with an underlying rate, the Poisson distribution describes their occurrences. *P*_0_(*τ*) is the zero-event probability density function resulting from the inhomogeneous Poisson process governed by the time-dependent event rate *z*_*µ*_(*τ*). Therefore the probability that no reactions (of any type, not just *µ*) occur in a time interval *τ* is given by the following equations:

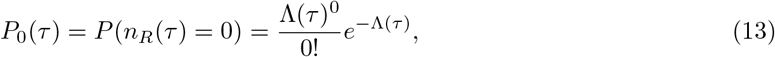

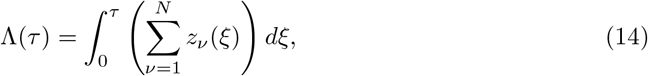

Where *n*_*R*_ is the number of reactions and *ξ* is a dummy time-variable. Combining (12), (13), and (14), we can now express *P*(*τ, µ*) in terms of the two parts described above:

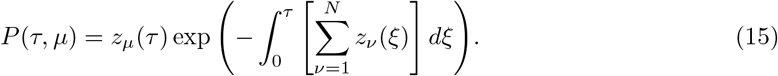

The probability that a specific *R*_*µ*_ reaction happens is a function of all other possible reactions due to the summation term in the exponential. For the next reaction to be of type *R*_*µ*_, the reactions *R*_*ν* ≠ *µ*_ and the specific reaction *R*_*µ*_ must not yet have occurred. The final step in computing *P*(*τ*) is to sum over all possible reaction types:

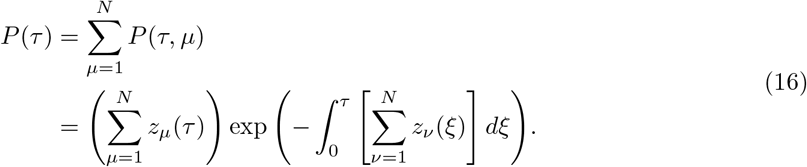

This is the probability density function describing the likelihood of the wait-time until the next reaction. To verify that this probability density function is properly normalized over its defined domain, we must show that 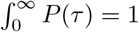. We recognize that (16) takes the following form:

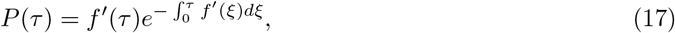

Where 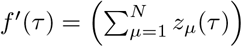. For the set of functions *f* where *f*(0) = 0 and *f*(*τ >* 0) integrating yields:

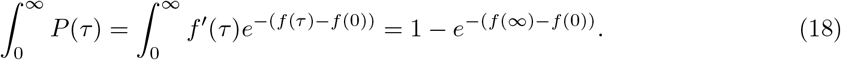

For the set of functions where *f* ′(*τ*) ≥ 0 and therefore *f*(*τ*) ≥ *f*(0), (18) is bounded with the following inequality:

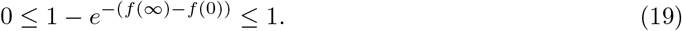

This is true here because for all *R*_*µ*_ reactions, *z*_*µ*_(*τ*) ≥ 0. The circumstances where 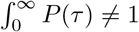 describe situations where there is a finite probability that no further reactions occur and provide insight on the reactive process: for a reaction to occur, *f*(∞) must not remain bounded and therefore *f* ′ must not decay to 0.

### 2.3. Gillespie Algorithm Equivalence

We are now at a point to ask if this reaction probability density function is equivalent to the Gillespie derivation in the limit of a fast diffusion rates relative to reaction rates. We begin by demonstrating the following equality:

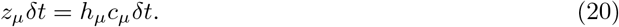

In a bounded system governed by the limit of fast diffusion, each molecule has equal probability of being anywhere within *V*, so the pseudo-probability distributions defined in (6) are sums of the uniform distribution over *V* :

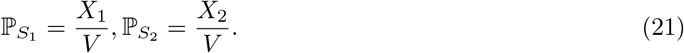

From reference [12], the right hand side of (20) is equal to:

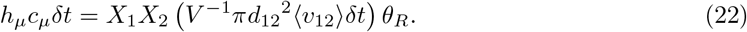

Substituting the values from (21) into (12) gives:

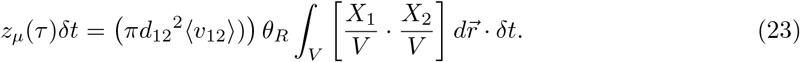

All the terms within the integral are constants with respect to position, therefore (23) reduces to

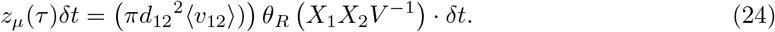

Which is exactly equal to *h*_*µ*_*c*_*µ*_*δt* in (22). To establish equivalence of *P*(*τ*) in this limit, we recognize that *z*_*µ*_ defined in (24) is time independent and the inhomogenous Poisson process used to derive (16) becomes a homogenous one:

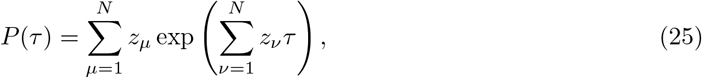

Which, using the demonstrated relationship in (20), is equivalent to *P*(*τ*) of the Gillespie algorithm [12].

### 2.4. Sampling 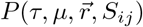 by decomposition of P(τ)

The joint probability distribution 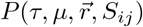 may be sampled through the use of conditional probabilities in a series of steps:

1. What is the wait time *τ*?
2. Given a wait time *τ*, what reaction *R*_*µ*_ took place?
3. Given a wait time *τ* and a specific reaction *R*_*µ*_, where did it take place?
4. Given a wait time *τ*, specific reaction *R*_*µ*_, and location 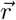, which molecules *S*_*ij*_ were involved?

These questions are the four terms on the right-hand side of the following equation and may be answered by sampling each distribution using Monte Carlo methods:

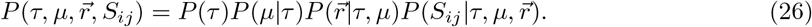

In this section, we proceed through the sampling of each distribution using a computational implementation for a simple bimolecular reaction as a guide (Fig 1A).

**Figure 1:**
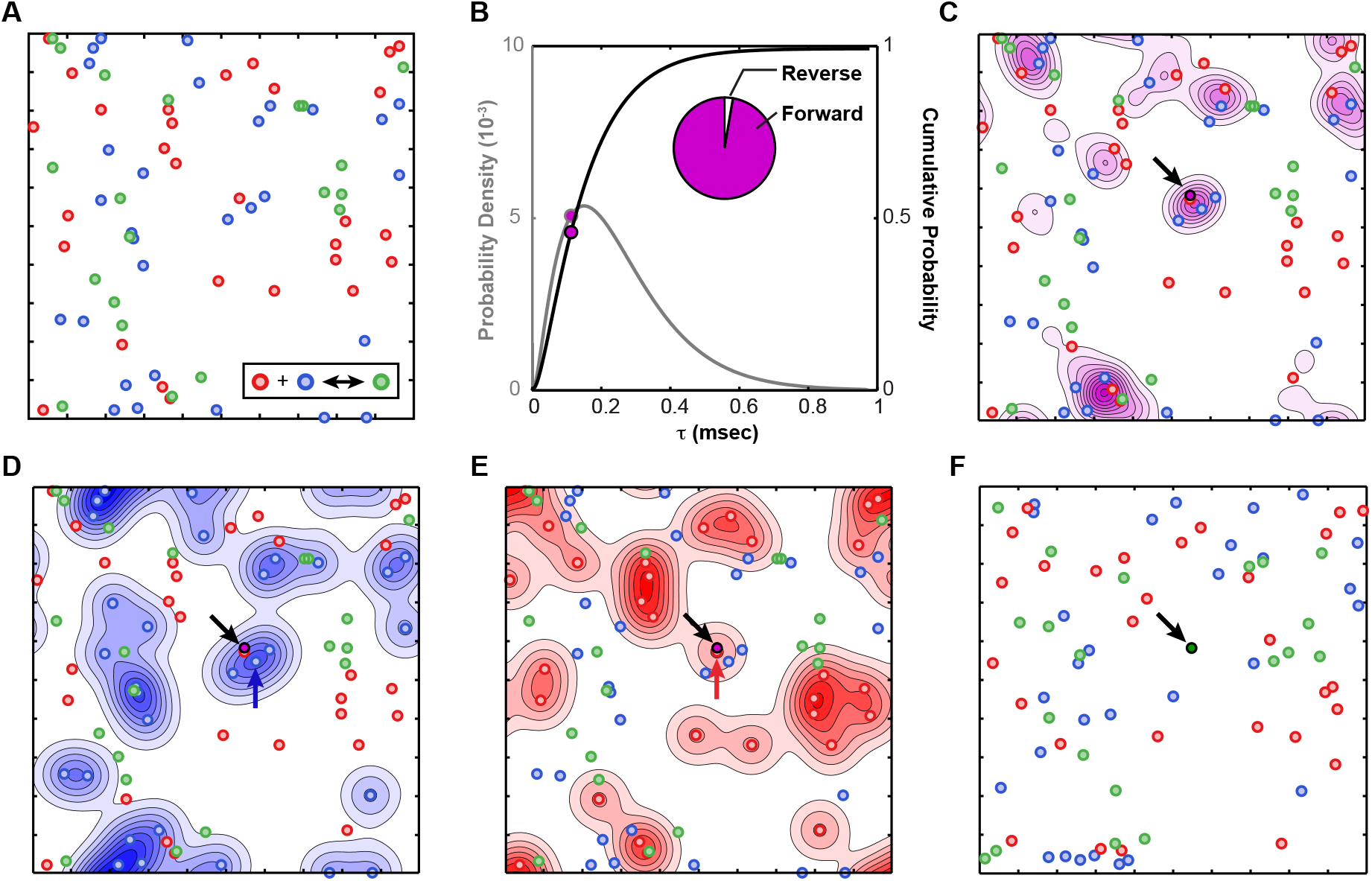
Simulation of a single reaction step. A. Initial positions of molecules. Molecules are randomly distributed over the environment. B. Reaction probability density function, *P*(*τ*) shown in gray and the cumulative probability is shown in black. The cumulative probability distribution is sampled via the inversion method, and the resulting value of *τ* is the wait time (magenta dots). The relative probabilities of the forward and reverse reactions occurring at this wait time is shown in the inset; random sampling selects the forward reaction. C. The spatial distribution of the reaction likelihood, 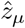, and the sampled reaction location (magenta dot). D, E. The location density functions 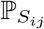 for *S*_1_ and *S*_2_ respectively. The *S*_1_ and *S*_2_ molecules likely to have participated are selected from these distributions. F. The updated system with removal of reacting *S*_1_ and *S*_2_, new *S*_3_, and displacement by diffusion of non-reacting molecules.

*What is the wait time τ ?* The reaction wait-time *τ* can be directly obtained by inversion sampling of the probability density function *P*(*τ*) shown in (16), providing the time at which the next reaction occurs (Fig 1B). Inversion sampling is a general method for random sampling of a probability distribution given its cumulative distribution function. Cumulative distribution functions are monotonic and map the domain of the probability distribution to a range between 0 to 1. Inversion sampling selects a number on the interval 0 to 1 from a uniform random distribution. The cumulative distribution function maps this value from the range back to a unique value of the domain.

*Given a wait time τ, what reaction R*_*µ*_ *took place?* The conditional probability *P*(*µ*|*τ*) can be expressed using (15) and (16) as:

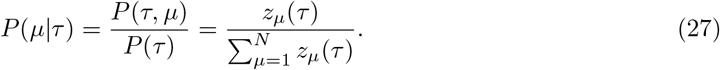

This is the the proportion of the total event probability that each reaction contributes. This is a discrete distribution with respect to the reaction variable *µ* and can be sampled to obtain the type of reaction that occurs (Fig 1B inset).

*Given a wait time τ and a specific reaction R*_*µ*_, *where did it take place?* The third question is answered by sampling the probability distribution 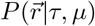, which represents the relative contribution of each point in space towards the occurrence of the reaction *R*_*µ*_ and is given by the following equation:

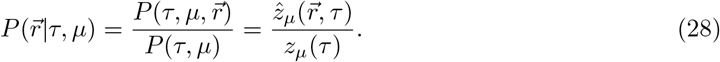

Additionally, because 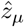 itself is a joint probability distribution over the number of spatial dimensions in the system, further decomposition needs to be done to sample each dimension (Fig 1C).

*Given wait time τ, specific reaction R*_*µ*_, *and location* 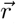, *which molecules S*_*ij*_ *were involved?* To answer the final question, we need to sample the probability distribution 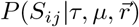. That is we need to answer the question: given a reaction, time and position, which of the involved molecules participated? This quantity can be expressed by the following:

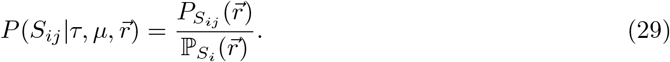

This is a discrete probability distribution where the probability of the *j*^*th*^ molecule being picked is the proportion of P it contributes at point 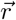. Sampling of this probability distribution is again performed using the inversion method as described in Question 2. This computation is performed for each species participating in the reaction (Fig 1D and E).

The reaction step has now been fully characterized. The reacting particles can be removed from the list of *S*_*ij*_ and the created particle can be added to *S*_*ij*_ at the location of the reaction, updating the quantities *X*_*i*_. The final step is to update the positions of the molecules that did not react this step but have moved due to diffusion, by sampling the individual probability density functions 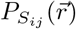. This completes one round of the simulation and leaves the system ready for the next (Fig 1F).

## 3. Results

### 3.1. Diffusion-Influenced Bimolecular Reactions

To verify that this algorithm produces expected behavior from reaction-limited to diffusion-limited regimes, we simulated a simple bimolecular association-dissociation reaction on a two dimensional domain. In two dimensions, effects due to local concentration differences as might arise in a reaction-diffusion system are exaggerated because the probability of two given molecules reacting is much more strongly dependent on separation distance than in three dimensions [11]. The bimolecular reaction is defined as 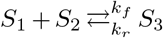, where *k*_*f*_ and *k*_*r*_ are the microscopic rate constants for association and dissociation, respectively. The algorithm was implemented using the MatLab programming language (Natick, MA).

This reaction was simulated across a range of values for *k*_*r*_*/k*_*f*_ and diffusion constants for species *S*_1_ and *S*_2_. Under conditions of fast diffusion where mixing occurred between reactions, the mean count of *S*_1_ closely matched the mean-field expected value Fig 2 (blue points). In the setting of slow diffusion, the effective association rate is expected to depart from the microscopic value due to the persistence of spatial correlations in molecular position created by the dissociation reaction. The dissociation of an *S*_3_ molecule produces an *S*_1_ molecule and an *S*_2_ molecule separated by only a short distance. These two molecules can either diffuse away from each other or re-associate. Under conditions of fast diffusion, re-association is less likely and the temporary spatial correlation between the two reactants is more quickly lost. Under conditions of slow diffusion, re-association frequently occurs, shifting the expected values for species counts. Indeed as the diffusion constants for *S*_1_ and *S*_2_ molecules were lowered, the observed *S*_1_ count shifted to the right, indicating that fewer *S*_1_ molecules were present than expected given the rate constants alone Fig 2 (red and yellow points).

**Figure 2:**
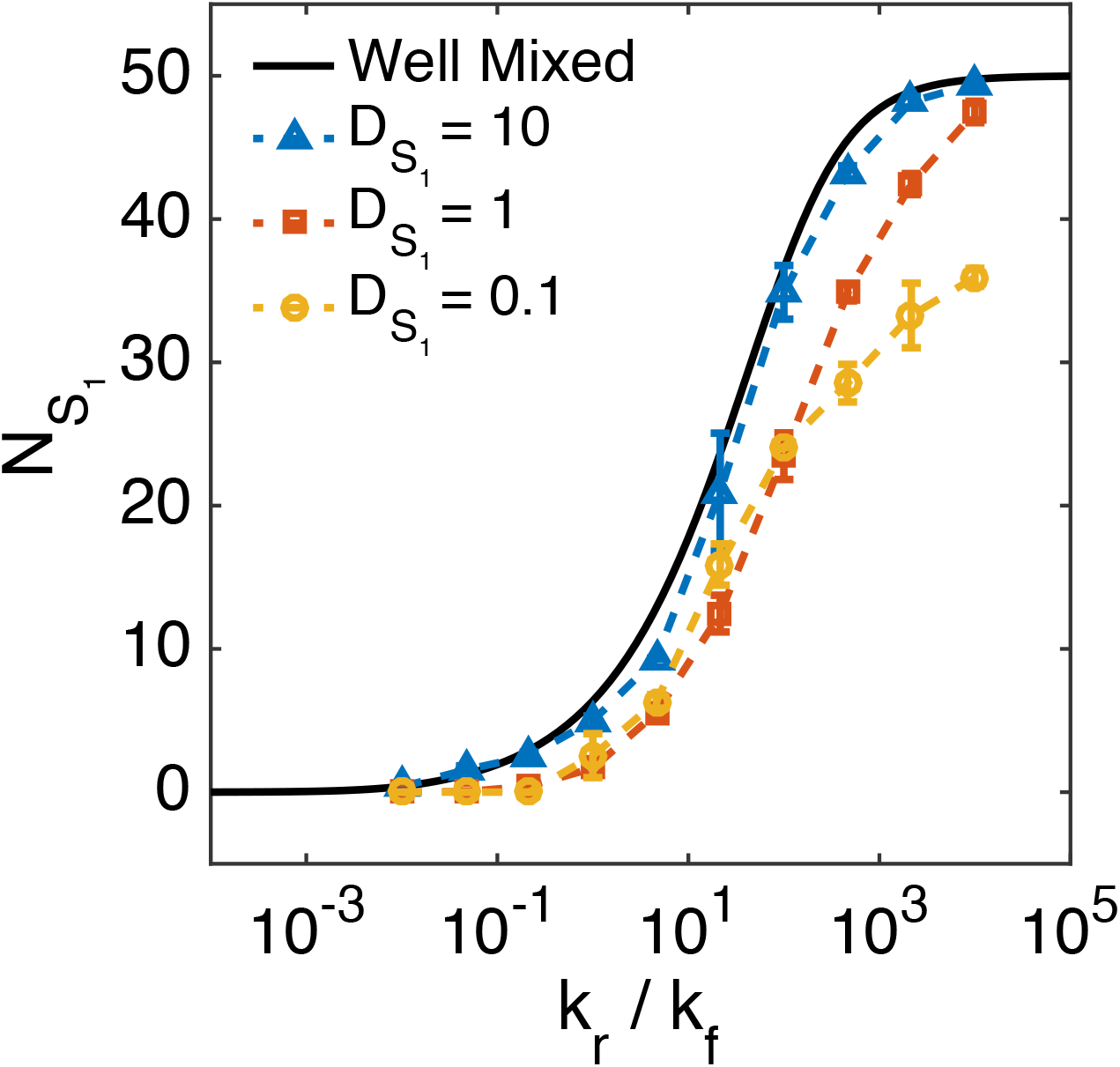
Association-dissociation reaction with re-binding effects. Expected number of *S*_1_ molecules computed under a mean-field approximation or simulated as a function of reactant diffusion constant. The results of the mean-field approach are shown in the black line, and simulation results are shown by the dashed lines and symbols. The diffusion constant for *S*_1_ ranged from 10*µ*m^2^s^−1^ to 0.1*µ*m^2^s^−1^. Additional parameter values used in the simulation are as follows: initial *S*_1_ = 30, initial *S*_2_ = 30, initial *S*_3_ = 20, 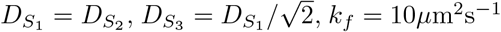 and *k*_*r*_ is set by *k*_*f*_ and the ratio given by the x-axis. The simulation size was 1*µ*m by 1*µ*m, solved on a 200 by 200 point numerical domain. 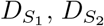, and 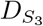 are diffusion coefficients of the species given by the subscripts.

### 3.2. Efficient Computational Strategies

A fundamental question for all reaction-diffusion simulations is how computational time scales as the number of molecules in the system increase. An ideal algorithm would have low start-up computational costs and minimal increases in computational time as the number of molecules increases. In situations where the number of molecules is large, this scaling property may be significantly more important than any start-up costs. The proposed algorithm has the potential to be computationally intensive, especially in regards to the generation of the pseudo-probability distributions 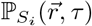, which requires the summation of individual probability distributions for each member of a species. However, though the use of specific computational strategies, the algorithm can be implemented in an efficient manner with highly desirable scaling in computational time as molecular number increases. This relies on the fact that 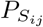 evolves identically for a single species except for a shift in position, providing an alternative strategy for computation employing convolution. For particles whose positions evolve according to a general probability distribution function 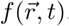, the function 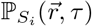 can be re-defined as:

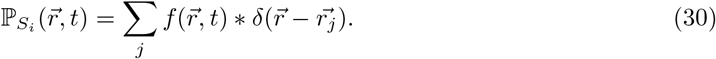

Where *δ* is the Dirac delta function. This convolution may be implemented directly or through use of Fourier transforms:

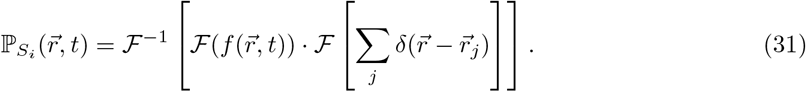

Where ℱ is the Fourier transform operator. In this approach, the summation of complex functions is avoided by converting calculations to a single multiplication step in frequency-space.

To demonstrate the computational efficiency of the convolution strategy, the time required to compute 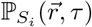 for an increasing number of particles was performed. The direct summation method was compared to direct convolution and Fourier transform based convolution where the total number of particles, *N*, ranged from 10 to 10^4^. While computation time scaled linearly with *N* for the direct sum method, computation time for both convolution based approaches was independent of particle number (Fig 3). This reflects optimum scaling that may or may not be realizable for all computational implementations of the simulation. It may be desirable to scale the numerical resolution with the particle number to maintain the same strength in the assumption underlying (6). In this case, the number of elements in the simulated domain scales with the number of particles. Therefore the efficiency becomes bounded by that of the Fast Fourier Transform algorithm at *N* log(*N*). Regardless of which approach is taken when increasing the number of particles within a simulation, a computational efficiency that scales for *N* particles between a lower bound of zero-order and upper bound of *N* log(*N*) is a highly advantageous feature of this algorithm.

**Figure 3:**
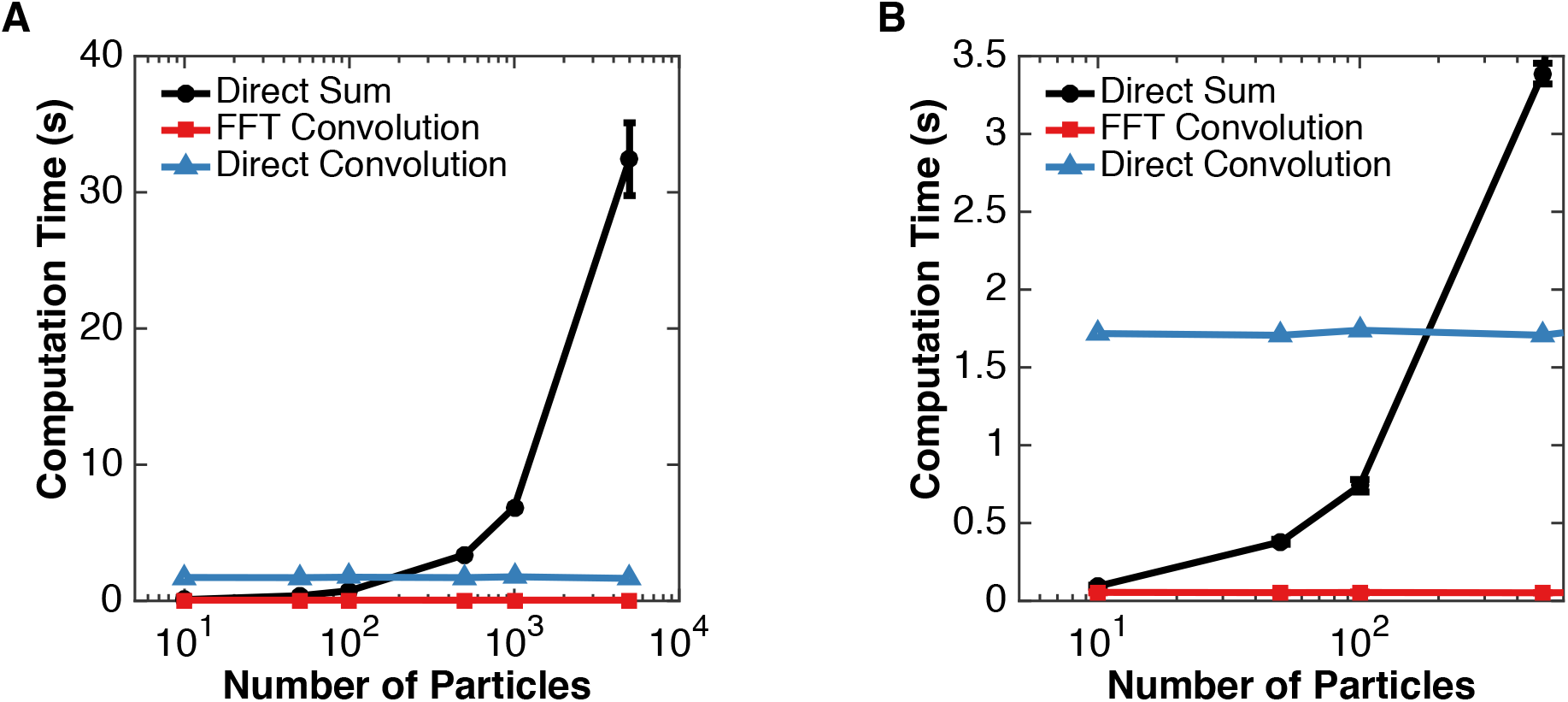
Comparison of computational efficiency for summation and convolution based approaches. A. Time to compute 𝕡_*S*_ as a function of number of *S* molecules for summation, direct convolution, and Fourier transform convolution. B. Magnification of cases between 10 and 500 molecules to highlight relative efficiencies for small numbers of molecules. Parameter values used for testing computational efficiency were: *D* = 1*µ*m^2^s^−1^, *τ* = 10^−3^s, where *D* is the diffusion coefficient and *τ* is as defined in the main text. The domain size was 1*µ*m by 1*µ*m discretized on a 1024 by 1024 grid.

### 3.3. Diffusive Mobility and Enzyme Efficiency

The formation of protein complexes is a ubiquitous part of biological systems [25, 26]. Much work has focused on understanding how conformational changes induced by protein-protein interactions can modulate intrinsic reaction rates through allostery [27]. These interactions also lead to the formation of macromolecular complexes or subcellular compartments where mobility of proteins differs from free diffusion. These structures, like lipid rafts and scaffolds, are known to impact cell signalling[28], but the direct effect of changes in mobility on reaction kinetics are less well understood. Using a molecular dynamics approach, Soula et. al show that changes in mobility can impact apparent binding affinities, but recognize computational challenges in using a molecular dynamics framework for modeling these processes [29]. The algorithm proposed here provides an efficient avenue to study these systems. In this section we examine a model reaction-diffusion system motivated by quantifying the effect that protein complex formation may have on enzymatic activity simply through the change in diffusivity that results from forming a macromolecular assembly. The effect of spatiotemporal correlations introduced by decreased diffusion rates or the restriction that comes from reactions confined to a 2-D surface is known to plan an important role in the kinetics of reacting systems [30, 31, 32, 33, 34]. Rebinding events which can occur with slow diffusion or poor mixing can lead to different macroscopic behavior than predicted from intrinsic microscopic rates [35, 36]. The simulation method introduced here provides a direct way to study these systems.

Consider a system where a substrate is produced in a spatially restricted manner and acted on by an enzyme that is capable of forming complexes with other proteins. The reaction scheme used in the simulation is shown in Fig 4A. A molecule S is produced by an essentially fixed point source and diffuses away subject to degradation, creating a spatial gradient in the probability of encountering S. An enzyme A binds S and converts it to P. Alternatively, while bound to S, A can be altered in a manner that leads to a change in mobility but does not affect intrinsic enzymatic efficiency. This scenario represents a generalization of a number of specific mechanisms that could lead to spatial localization of an enzyme near a source of substrate. This may reflect a change in phosphorylation state or conformation that leads to protein complex formation, increased lipid-bilayer binding, or any alteration that reduces the diffusion rate of A. As a result, the mobility of A is reduced in a region where S is likely to be found.

**Figure 4:**
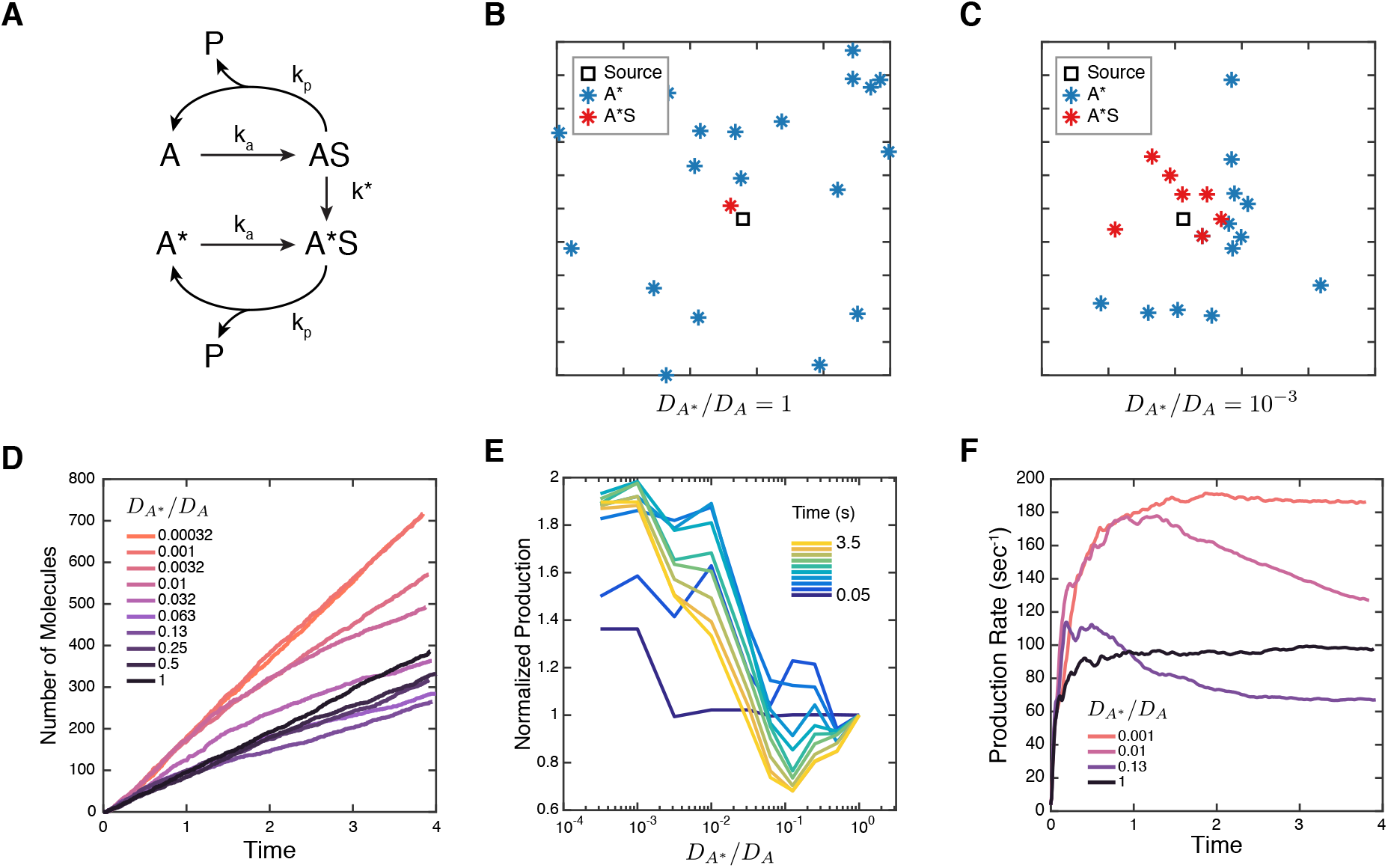
Changes to enzyme mobility alter effective production rates. A. Diagram of reaction. Substrate S is produced by a relatively stationary point source. Enzyme A binds this substrate and converts it to a product P. Alternatively, when bound to the substrate, A can be converted to a form with lower mobility, A* with identical intrinsic catalytic ability to turn S to P. B. Snapshot of enzyme positions and occupancy after 8000 reaction time-steps with no reduction in mobility. The source of S is denoted by the black square. C. Equivalent to B but with a thousand-fold reduction in mobility. Enzymes are clustered near the substrate source and are more likely to be bound to S. D. Total number of P molecules produced as a function of time for a range of reductions in diffusivity. E. Equivalent data to D showing total production as a function of diffusion constant reduction at different times. For intermediate reductions in diffusion, production initially exceeded but then fell below the control case. F. Production rate for selected reductions in mobility spanning three orders of magnitude, computed by a sliding linear-regression to find the slope of curves in D. For high reductions in mobility, production rate doubles the control case. For lesser reductions, production rate initially exceeds the control case but diminishes with time. Parameter values used in the simulation were: *D*_*S*_ = *D*_*A*_ = 1*µ*m^2^sec^−1^, substrate production rate = 1000 sec^−1^, substrate decay rate = 300 s^−1^, *k*_*a*_ = 10*µ*m^2^s^−1^, *k*^*∗*^ = 100*µ*m^2^s^−1^, *k*_*p*_ = 20*µ*m^2^s^−1^. The domain size was 1*µ*m by 1*µ*m discretized on a 200 by 200 grid. Here *D* variables denote diffusion coefficients and *k* variables denote reaction rates.

This system provides a generalized scenario for examining the effect that protein complex formation has on enzymatic efficiency even if intrinsic catalytic rates remain constant. Figure 4B and C show a snapshot of the enzyme positions at the conclusion of 8000 reaction time-steps for the case of no mobility reduction and a 1000-fold reduction. As expected, enzymes where mobility reduction is high are concentrated in the vicinity of the source of S, indicating successful recruitment near the source of substrate due to the reduction in mobility. The production of P depends on the relative mobility of *A*\* and is higher in cases where reduction is the greatest but changes as the system evolves in time (Fig 4D-F). For large reductions in mobility, the production rate of P is higher than if no reduction occurs and remains so for thousands of reaction steps. In this case, although the enzymes search space less quickly, they remain in a region with an increased probability of encountering S, leading to higher overall efficiency. For intermediate drops in mobility, however, the production rate of P initially exceeds but eventually falls below the control level. Here, diffusion is not reduced enough to maintain the high degree of initial spatial correlation with the source of S to balance the effect on search-times.

## 4. Discussion

In this work, we use probabilistic techniques to simulate a general reaction system in both space and time. The approach is driven by reaction events, is capable of spanning diffusion limited to well-mixed regimes, and is shown to be mathematically identical to the Gillespie algorithm in the limit of rapid diffusion. Using a computational implementation of this algorithm, we demonstrate that it captures equilibrium values in well-mixed systems and re-binding events when diffusion is slow. Lastly, in a model of protein complex formation, we find tradeoffs may exist between spatial localization and search-time due to macromolecular assembly. For simplicity, this algorithm will be referred to as the Spatiotemporal Gillespie algorithm (STG).

The STG method relies on probabilistically jumping between events in order to skip details that do not change the progression of the system in a manner similar to GFRD and FPKMC methods [16, 20, 17]. The primary difference between these methods and STG is the definition of an event and the construction of the underlying wait time distribution. GFRD and FPKMC are event driven, but the event definition is related to probability of multi-way-interactions rather than successful reactions. In the regime of highly diffusion-limited reactions, this is equivalent, but when interactions do not guarantee reactions, these definitions differ. In this case, both GFRD and FPKMC increment through time by simulating many unsuccessful interaction events before a reaction occurs. By focusing on reactive events, the STG algorithm avoids this potential inefficiency that may arise when a near-diffusion-limited process or even a reaction-limited process is part of the system. As a result, the STG algorithm can simultaneously simulate reaction-limited (highlighted by its Gillespie algorithm equivalence) and diffusion-limited processes that share components. This hybrid functionality is a fundamental strength of the STG algorithm.

To achieve this, the STG algorithm relies on fundamental assumptions in both the analytical definition and computational implementation. The primary assumption of the analytical definition is that non-reactive collisions with labeled particles (those that have been given an identity in the system) do not significantly alter the random-walk statistics that arise from collisions with all other unlabeled particles in the environment. This assumption may not hold if electrostatic interactions are involved that disrupt diffusive motion [37]. This is a challenge not unique to this approach, but may be handled here by defining the diffusion-altering event as a separate reaction in the system and creating an intermediate species for more precise treatment.

The primary assumption in the numerical implementation of the STG algorithm is in the numerical resolution used for the pseudo-probability distribution ℙ_*S*_, which approximates the probability that a single molecule resides at a position. In theory, collision volumes are vanishingly small and the probability that two molecules simultaneously fall within it approaches zero. In this limit, the union probability is well approximated by summing individual probabilities and neglecting the intersection probabilities. In practice, the minimum value that individual probabilities may take depends on the resolution chosen for numerical approximation of the individual probability density functions. Therefore, to maintain the accuracy of this assumption, the numerical resolution must be kept high enough that the probability of a single particle of a given type residing in a numerical discretization of space is small such that the probability of two remains negligible. This approach, however, makes the STG approach highly flexible. Any motion that can be probabilistically described can be incorporated into the framework, allowing for the study of systems where body forces play a role and processes in 1, 2, and 3 dimensions. Furthermore, systems with mixed dimensionalities, for example a 1D DNA strand within a 3D environment, could be modeled with the STG framework.

The analysis on computational efficiency demonstrates that the algorithm exhibits efficient scaling as the number of particles increases when convolution-based methods are employed. In the best-case scenario, when the resolution of numerical implementation does not need to be changed, scaling is zero-order: additional particles do not effect the computational time for a given reaction. However, even if the numerical resolution is scaled with the number of particles, the computational complexity follows that of the method chosen for convolution. While the computations required to implement the STG algorithm are more involved than simulating diffusive trajectories or subvolume transitions, its reaction-event driven nature and scaling properties are highly advantageous features as parallelized CPU and GPU computing power continues to grow.

The simple model of protein complex formation presented here demonstrates two fundamental competing factors in reaction diffusion systems: control of protein localization and effects on search times. In all cases, the enzymes A are likely to undergo a reduction in mobility where S molecules are more prevalent and do so with the same kinetics. Such a change may occur when substrate binding influences covalent modification of the enzyme itself, leading to changes in binding partner affinity as is seen in Rho GTPase enzymes [38]. This localization in a zone of higher levels of S leads to increased production rates. As time progresses, enzymes that only underwent a moderate reduction in mobility begin to spread out and production drops. These enzymes are slower to search space and are no longer restricted to a region with high local concentrations of S and as a result, production falls below that of the case where no reduction in mobility occurs. In summary, protein complex formation increases effective enzyme efficiency through localization near a source of substrate. However, maintenance of spatial correlation with the source becomes of increased importance to avoid the disadvantageous lengthening of search-times that results from mobility reduction.

The methods proposed here provide a rigorous framework for assessing the effects of slow diffusion and small numbers of molecular species as well as the conditions where these considerations become important. Furthermore, this unique approach unifies reaction-limited and diffusion-limited regimes within a single framework, allowing of straightforward simulation of biological systems where reaction of both regimes are coupled. We anticipate our framework will help decipher the complex interplay between reaction and diffusion in biological signaling.

## 5. Acknowledgments

We would like to greatly thank Elliot Elson for his input and many useful discussions. We would also like to acknowledge Mauricio Del Razo Sarmina and Hong Qian for their valuable and constructive feedback. This work was supported by the National Institutes of Health Grants T32 EB018266 and T32 GM007200 to A.J.L.

